# Deep learning reveals hidden variables underlying NF-κB activation in single cells

**DOI:** 10.1101/687848

**Authors:** Parthiv Patel, Nir Drayman, Ping Liu, Mustafa Bilgic, Savaş Tay

## Abstract

Individual cells show great heterogeneity when responding to environmental cues. For example, under cytokine stimulation some cells activate immune signaling pathways while others completely ignore the signal. The underlying sources of cellular variability have been inaccessible due to the destructive nature of experiments. Here we apply deep learning, live-cell analysis, and mechanistic modeling to uncover hidden variables controlling NF-κB activation in single-cells. Our computer-vision algorithm accurately predicts cells that will respond to pro-inflammatory TNF stimulation and shows that single-cell activation is pre-determined by minute amounts of “leaky” nuclear NF-κB localization before stimulation. Theoretical analysis predicts and experiments confirm that the ratio of NF-κB to its inhibitor IκB determines the activation probability of a given cell. Our results demonstrate how computer vision can study living-cells without the use of destructive measurements and settles the question of whether heterogenous NF-κB activation is controlled by pre-existing deterministic variables or purely stochastic ones.

---

Individual cells show unpredictable and highly variable responses in a wide range of contexts, from immune signaling to drug response^1–5^. For example, following stimulation with signaling molecules like tumor necrosis factor (TNF) and lipopolysaccharides (LPS), a portion of cells in a population will activate inflammatory response pathways like NF-κB while others will completely ignore the stimulus^6^. Determining the sources of signaling variability is of immense importance for fundamentally understanding gene regulation, signaling, immunity, and development, as well as in predicting variable drug responses and tolerance^7–10^. Despite previous characterization of heterogeneous NF-κB responses in a wide range of contexts^6,11,12^, it remains extremely difficult to explore the molecular and cellular mechanisms that drive variable behavior in single cells.

NF-κB is a key transcriptional pathway that is activated by a plethora of signaling molecules^13^ and controls the expression of hundreds of pro-inflammatory and cell fate genes^14^. Dysregulation of NF-κB is implicated in many physiological conditions including infection, autoimmunity, and cancer^13,15^. Live-cell analysis has shown that NF-κB activation is highly variable in single cells, where many cells show complete cytoplasm-to-nucleus translocation of the p65 transcription factor (the hallmark of pathway activation), while others ignore the stimulus and show no translocation and no NF-κB target gene expression (Figure 1a). While the fraction of cells that respond to a stimulus increases in a dose-dependent manner^6^, it is unclear whether these cell-to-cell differences are due to stochastic, random processes or are pre-determined by variability in unknown molecular components that are hidden to researchers (Figure 1b). Despite this seemingly noisy behavior, NF-κB nevertheless manages to mount very specific responses to different signaling molecules, taking into account their concentrations and temporal ordering^1,6,11^.

**Figure 1:**
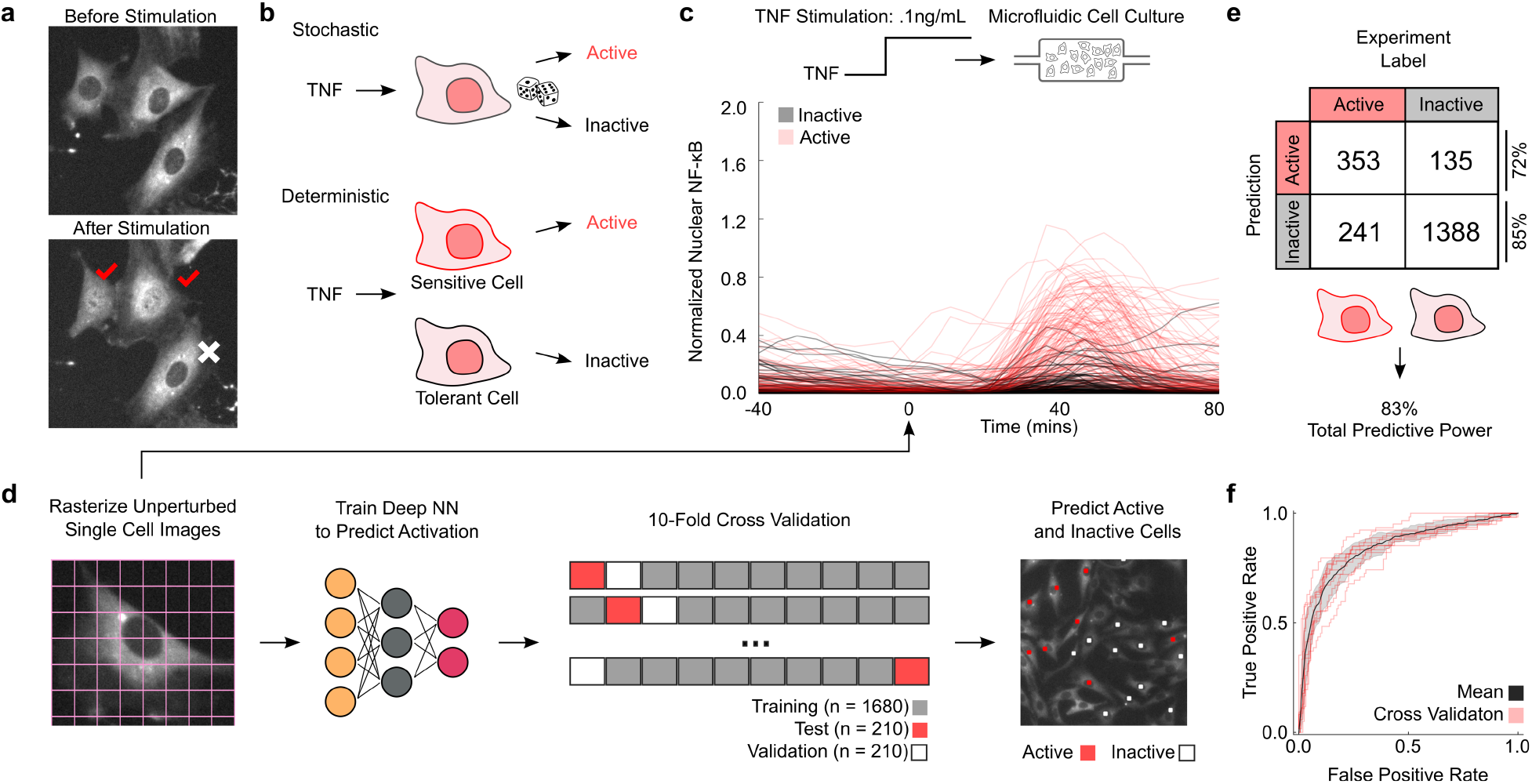
Deep learning predicts NF-κB activation in single cells. This computational prediction approach allows experimentally accessing unperturbed cell states to investigate the underlying mechanisms in variable NF-κB activation. **(a)** Under stimulation with TNF, a fraction of individual cells persist in an un-activated state and do not show nuclear NF-κB localization, while others activate and NF-κB transcription factor p65 accumulates in the nucleus. Example images show activated cells, indicated with red check marks. **(b)** Analysis of single cells under constant TNF dose shows variable single cell activation in the population: a given cell may or may not respond to the TNF stimulus. It is unclear whether single cell variability is due to purely stochastic processes (i.e. if a given cell can randomly achieve activated state), or if it is deterministic where only sensitive cells activate under TNF input. **(c)** We use microfluidic cell culture to stimulate cells with TNF and image the nuclear localization of NF-κB over time in single cells. Analysis of individual cells reveals NF-κB localization traces (0.1ng/mL TNF stimulation shown on the right, stimulation starts at t=0). Single cell traces show heterogenous activation and subpopulations of active and inactive cells. **(d)** We record pre-stimulation images of mouse 3T3 fibroblasts that express p65-dsRed and H2B-GFP reporters and feed them into our deep learning pipeline, using 10-fold cross validation, to predict the identity of active and inactive cells after TNF stimulation. **(e)** Deep learning prediction of active and inactive cells from the full population yields 83% positive prediction. **(f)** The model significantly predicts activated and inactivated cells and has an ROC AUC of 0.753.

The challenge in understanding the cause of heterogenous responses in signaling and gene regulation is a classic uncertainty principle problem – we can only identify the responding (i.e. activated) cells after we chemically stimulate the population with signaling molecules, which will inevitably perturb the cellular states we wish to examine. Many signaling pathways, including NF-κB, contain multiple feedbacks that upon exposure to an external signal rapidly change the molecular composition of the pathway. This presents a fundamental difficulty in determining how cell-to-cell differences impact the probability of any given cell to respond to a stimulus. One way to circumvent this problem is by use of mathematical modeling^16,17^. Stochastic modeling of pathway dynamics can reveal important insights and general patterns of noisy events, but, are bound by the underlying assumptions of the models and can offer many plausible explanations for cellular heterogeneity. Because of these experimental and theoretical limitations, molecular mechanisms underlying important cellular behaviors like variable drug response, digital pathway activation, and signal tolerance currently remain unknown^6,18–20^.

To understand to what degree NF-κB activation is driven by unpredictable stochastic molecular fluctuations, or whether there are deterministic cell state drivers of NF-κB response, we adopted an image-based machine learning approach to predict which individual cells will activate the NF-κB pathway in response to an inflammatory stimulus. By live-imaging the cells before, during and after stimulation, we were able to use the cell image **before** stimulation to predict whether the cell will activate the NF-κB pathway or not. We developed an image based convolutional neural network (CNN) deep learning model aided by a support vector machine (SVM) to predict outcomes of chemical stimulation in individual cells (see **Supplemental Info**). Our computer-vision based method classifies cells into responding and non-responding groups based solely on the unperturbed cell’s image and is able to predict which cells will respond to or ignore TNF stimulation with 82±3% accuracy. As this prediction is done on unstimulated cells, our approach allows studying how the cell’s unperturbed molecular state differs between responding vs. nonresponding cells, which was not possible before this study. Here we applied this method to understand if and how the dynamics of NF-κB response is bound by non-stochastic factors.

To develop a predictive machine learning model for NF-κB activation in individual cells, we first performed experiments to generate a reference data set. We used a high throughput microfluidic cell culture platform^21^ to chemically stimulate and quantitatively measure NF-κB response in cultured mouse fibroblast cells. These cells express p65-dsRed and H2B-GFP fluorescent reporters^6^ to track NF-κB nuclear translocation in real-time. Cells were first imaged unperturbed for 1.5 hours, stimulated (using the automated microfluidic system) with TNF (0.1 ng/mL) and monitored for 6 hours. Custom image analysis software was used to track the nuclear localization of p65 and assign a label to each cell (activated vs. not-activated)^21^ (Figure 1c).

We then trained and tested the CNN deep learning algorithm on our annotated set of images of single cells (n = 2113), taken before stimulation with TNF (t=0). We used the three image channels (phase-contrast, nuclear marker [GFP] and the basal state of p65 [dsRed]) as inputs to the classifier. We performed 10-fold cross-validation with test and validation sample sets to computationally validate our predictions and made predictions on unstimulated cells (Figure 1d). In total the algorithm correctly classified the future activation state of 82±3% of the cells (the mean and s.e. of individual inference on subsets) (Figure 1e). The CNN schematic is shown in Supplemental Figure 1a and ROC curve is shown in Figure 1f. We can accurately predict individual cells response from their pre-stimulation images which strongly suggests the existence of deterministic causes underlying NF-κB response in single cells.

While highly powerful in predicting, CNNs are difficult to biologically interpret as they rely on pixel-level information rather than descriptive predictors. Thus, to gain further biological insight into the deterministic nature of NF-κB sensitivity, we used a second machine-learning approach, support vector machine (SVM) (**See Supplementary Info**), which relies on 236 defined predictors (Figure 2a). We also expanded the experimental dataset from a single TNF dose to a range of doses (0.005 to 5 ng/mL, n=3456, Supplementary Figure 2a). We identified a subset of highly predictive image features that correlate with the cell’s likelihood to become activated, including the basal (pre-stimulus) level and standard deviation of nuclear NF-κB and several texture features (Figure 2c, d). Using these features, cells can be visualized by t-stochastic neighbors embedding (t-SNE), which match “TNF-sensitive” and “TNF-resistant” cells clusters (Figure 2b).

**Figure 2:**
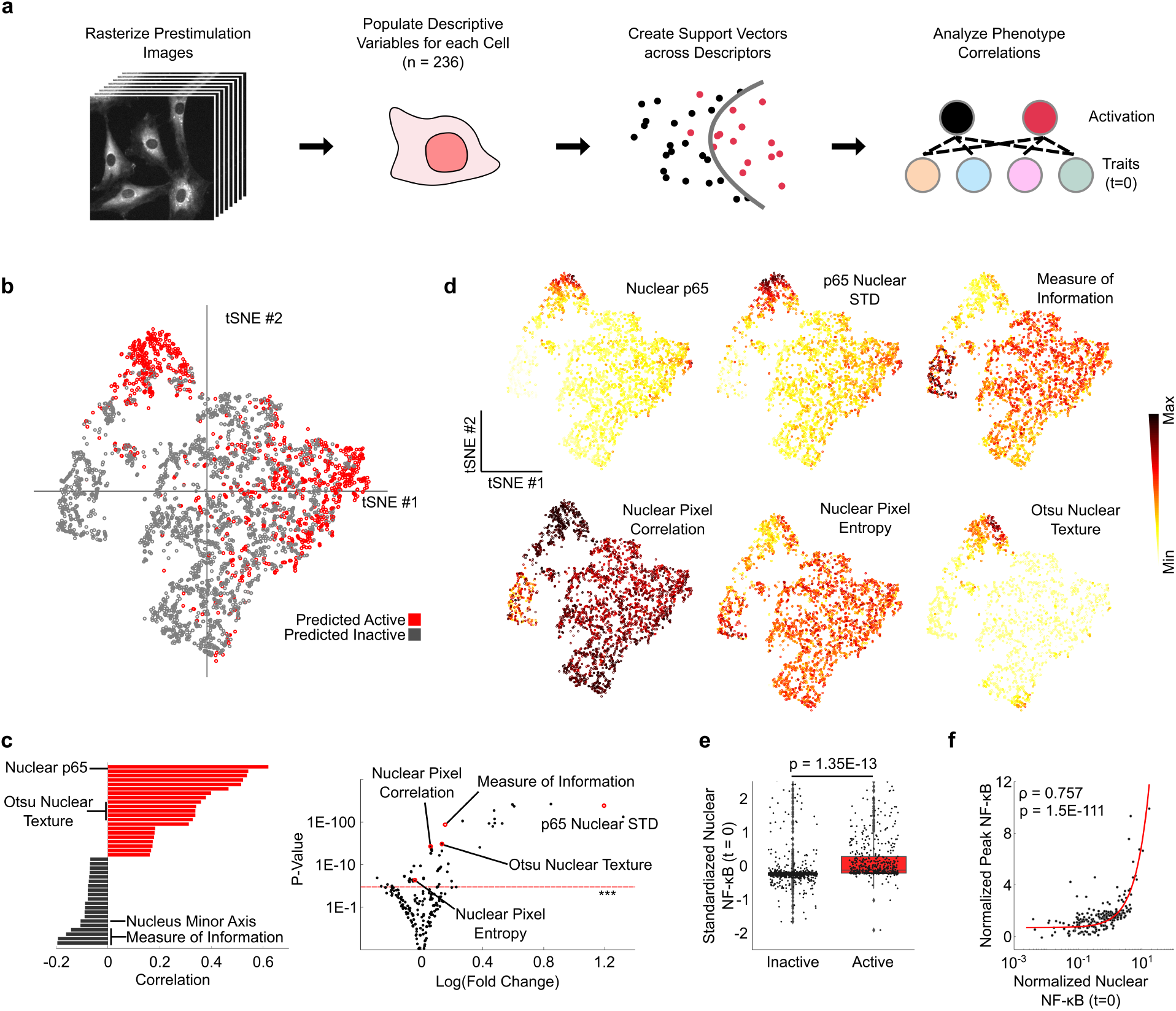
Support vector machine learning identifies “leaky” nuclear NF-κB localization as the primary predictor of NF-κB activation. **(a)** We feed pre-stimulation images into our SVM pipeline which identifies descriptive variables for the images and creates support vectors across the features allowing a connection between pre-stimulation traits and activation probability. **(b)** Using a subset of highly predictive features (n=8), tSNE corroborates the clustering of a highly predictive fraction of cells that are TNF sensitive and TNF resistant. **(c)** We determined the top feature of activation probability as mean nuclear fluorescence of the p65-dsRed signal in the nucleus before any exposure to TNF (i.e. leaky localization). Other highly predictive features include the standard deviation of nuclear p65, mean nuclear phase intensity, major axis length and a texture feature describing information measure of correlation in the nucleus, as well as aggregative image features like Otsu dimension, and SFTA (*** = 0.001). **(d)** tSNE plots of several highly predictive features align with predictions from SVM. **(e)** Pre-stimulation NF-κB nuclear fluorescence accounts for a high degree of variance and activated cells have a significantly higher level than inactive cells. **(f)** Activated cells show significant correlation between nuclear NF-κB at t=0 and NF-κB peak height after stimulation.

Further analysis of the contribution of individual image features to the prediction revealed that most of the variation in single cell predictions is explained by a single feature – the nuclear p65-dsRed levels before TNF stimulation (r = .62), which showed significant difference between TNF-sensitive and TNF-resistant cells. Cells that responded to TNF stimulus had, on average, a five-fold higher level of “leaky” nuclear p65 before stimulation than those that did not respond (Figure 2e, p = 1.35E-13), and nuclear NF-κB (p65) at t=0 was significantly correlated with the NF-κB peak height after stimulation (Figure 2f, p=1.5e-111). It is important to note that the low level of nuclear leakiness we observe is only about 12% of the total fluorescence in a given cell and is far below what is seen during activation. This small but significant difference, which evaded detection until this study, shows that the regulation of the steady state (i.e. pre-stimulus) NF-κB localization is an important determinant of NF-κB activation and demonstrates the power of utilizing computer vision to analyze single cell responses.

There are many different regulators that can influence p65 nuclear localization in resting (i.e. unstimulated) cells. The NF-κB pathway is robust to environmental fluctuations and noise, and many built-in negative feedback mechanisms (Figure 3a) prevent spontaneous activation (nuclear import of p65) in the absence of biochemical stimulation. Nevertheless, our imaging data clearly reveals that many cells show “leaky” nuclear p65 (localization without stimulus), and that this small but significant pre-stimulus p65 localization pre-determines the sensitivity of the cells in responding to upcoming TNF challenges.

**Figure 3:**
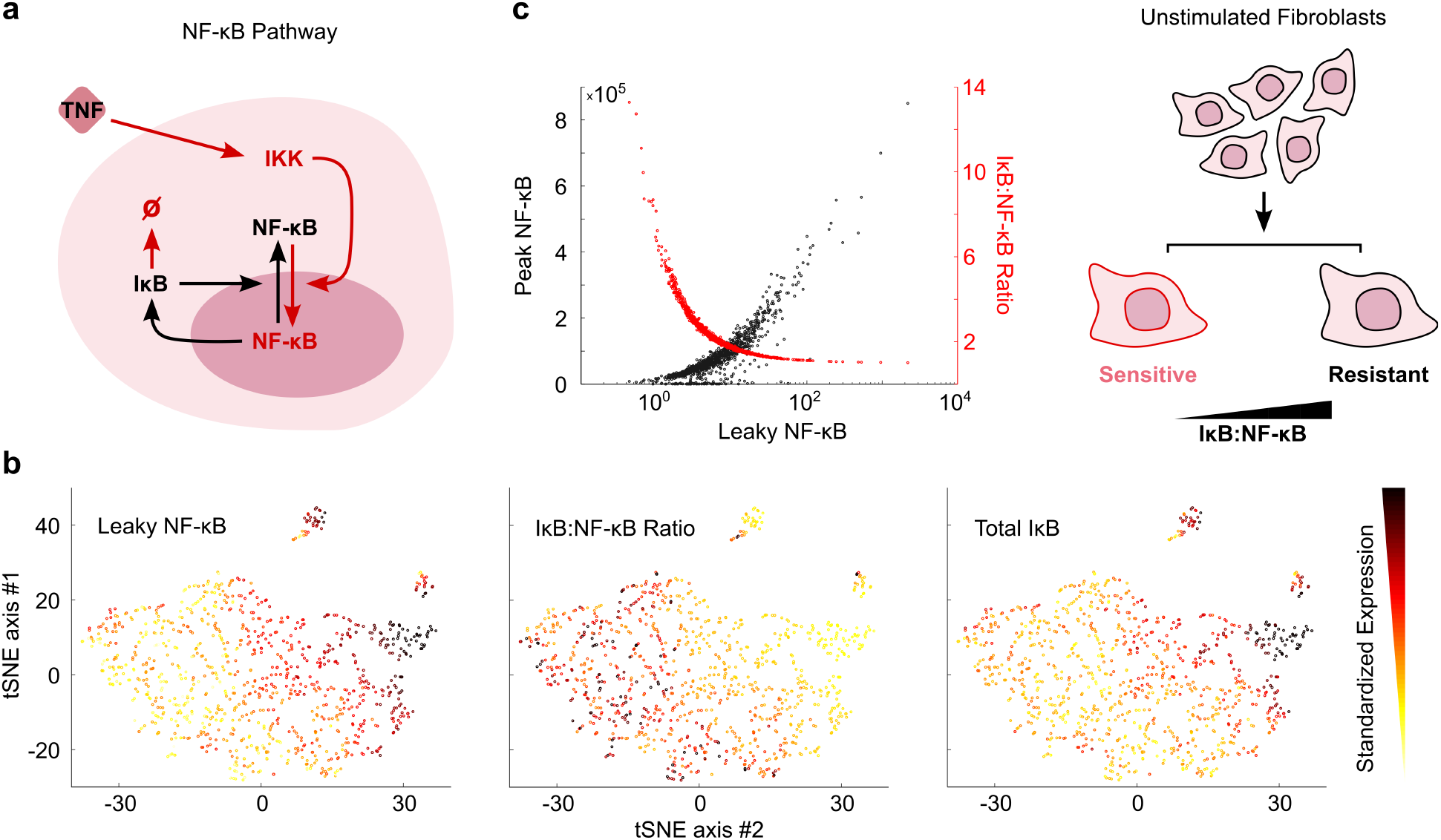
Mechanistic simulations suggest that leaky NF-κB localization and overall NF-κB activation is predetermined by the ratio of IκB to NF-κB proteins in single cells. **(a)** Simplified schematic of the NF-κB pathway used in simulations. IκB provides negative feedback to the pathway, preventing NF-κB nuclear localization in unstimulated cells. **(b)** tSNE mapping of simulated single cells with different NF-κB pathway protein levels shows that nuclear NF-κB level and IκB/NF-κB ratio are anti-correlated. **(c)** Simulations indicate that increasing IκB/NF-κB ratio make cells more resistant to activation under TNF stimulation.

To understand the molecular mechanism behind p65 nuclear leakiness and how it could lead to increased TNF sensitivity and NF-κB activation probability, we explored the mechanistic mathematical model of NF-κB pathway in single cells^6,12^. We found that variability in IκB levels, the main inhibitor of NF-κB that keeps it in the cytoplasm, could explain the observed NF-κB leakiness. In addition to binding to p65 and keeping it in the cytoplasm in unstimulated cells, IκB acts as a dynamic negative feedback regulator of the pathway, since IκB is a direct target gene of NF-κB. IκB is produced when NF-κB is activated and enters the nucleus. Using dynamic simulations, we perturbed the IκB/NF-κB ratio in single cells prior to TNF stimulation to determine the resulting likelihood of pathway activation for single cells (n=1000) (Supplementary Figure 3a). We find a major difference in cellular activation probability and peak height for different IκB levels (Supplementary Figure 3b). Counterintuitively, cells with high initial IκB levels require a smaller TNF dose to achieve NF-κB activation, whereas cells with low initial IκB levels are very resistant to any TNF dose. This surprising finding is explained by the facts that the probability of activation depends on the IκB/NF-κB ratio, and not the total level of IκB, and that the IκB/NF-κB ratio is anti-correlated to the total IκB level (Supplementary Figure 3b). This finding indicates that a pre-existing imbalance in the NF-κB negative feedback is responsible for increased TNF sensitivity, and that the activation probability of individual cells is pre-determined by the molecular ratio of IκB to NF-κB in the cell (Figure 3c).

To experimentally validate this prediction from mathematical modeling, we fixed and stained unstimulated cells for IκB-α protein expression and analyzed the relationship between IκB and NF-κB in individual cells using immunofluorescence (Figure 4a). Our deep-learning approach allows accurate prediction of the TNF stimulation outcome of these cells without actually exposing them to TNF, hence allowing the analysis of their unperturbed IκB protein states by immunofluorescence. We segmented the nuclear and cytoplasmic compartments and found that, as predicted by our simulations, there is a significant inverse correlation between the IκB/NF-κB ratio and leaky nuclear p65 localization (Figure 4b, ρ=−0.26, p=2.7E-11), which is the main feature that determines cell activation upon TNF stimulation.

**Figure 4:**
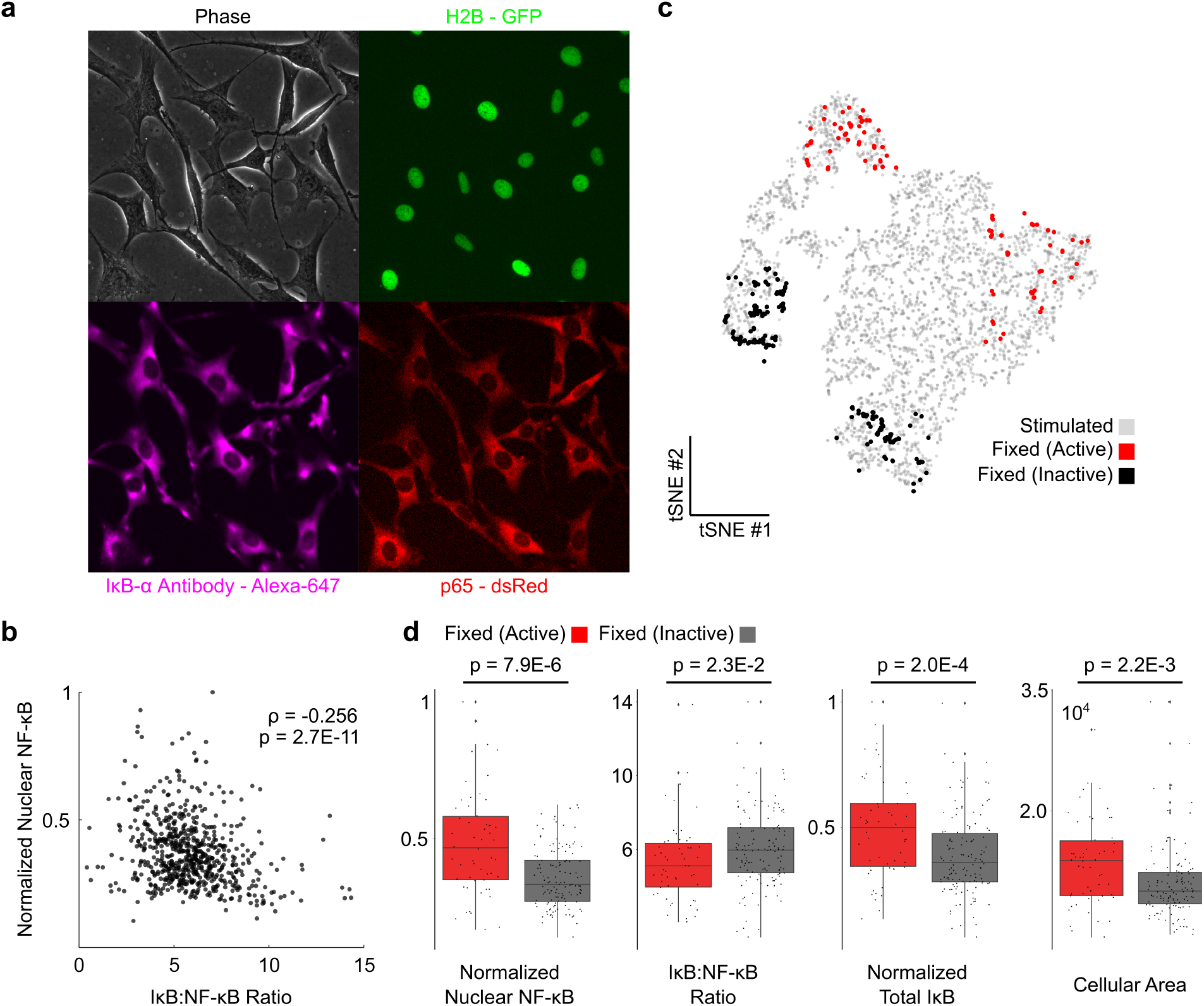
NF-κB activation is pre-determined by the NF-κB/IκB ratio in single cells. We validated our machine learning and modeling predictions by immunofluorescence staining experiments. **(a)** Image of unperturbed 3T3 cells fixed and stained with IκB-α with fluorescent p65 and H2B reporters. **(b)** There is a significant inverse correlation between leaky nuclear NF-κB localization and IκB/NF-κB ratio. **(c)** High scoring fixed cells are mapped onto tSNE. **(d)** IκB/NF-κB ratio is significantly correlated with nuclear NF-κB state at t=0 in predicted fixed cells, and there is a significant difference between IκB/NF-κB ratio, leaky nuclear NF-κB localization and total IκB as well as cellular area.

Next, we mapped the fixed-and-stained cells onto our tSNE visualization of the previous live imaging data (Supplementary Figure 2b) allowing us to infer IκB localization on our previous experiments for high scoring cells for activation (Figure 4c). We find that there is indeed a significant difference in nuclear NF-κB, IκB/NF-κB Ratio, and total IκB as well as cellular area, validating our prediction that IκB/NF-κB ratio is driving activation probability in NF-κB (Figure 4d).

In summary, our results show how image-based machine leaning can be used to study how cellular states affect the probability of seemingly stochastic events in signaling, as well as reveal the molecular determinants of these states. We found that the cell-to-cell variability in NF-κB activation is largely explained by a pre-existing difference in the ratio of the NF-κB and its inhibitor, IκB. It remains an open question as to what leads to this variation in the IκB/NF-κB ratio: one intriguing possibility is that epigenetic variance in genes encoding for NF-κB network components enforce the variable activation chance we observe. More broadly, the contrived variance we find in steady state IκB and NF-κB levels demonstrates the importance and potential functionality of gene expression variance in signaling. Overall, our demonstration of machine learning in the identification and elaboration of highly heterogenous signaling event leads from a theoretical prediction to an experimental observation of how cells could exploit heterogeneity as a non-trivial signaling driver and offers a novel tool to look at living cells through a prospective lens.

## Author Contributions

P.P. did microfluidic experiments; P.L. and P.P. ran machine learning pipeline; P.P. analyzed the experimental data with assistance from N.D.; M.B and S.T supervised the work.

## Correspondence

Savaş Tay, Institute for Molecular Engineering, The University of Chicago

## MATERIALS AND METHODS

### Microfluidic Cell Culture

Cell culture chambers are made of PDMS and coated with fibronectin (FC010-10MG) overnight. Cells are seeded at a constant density of ~20,000 cells/cm^2^. Cells are taken at 100% confluence by incubating with .25% trypsin-EDTA (25200-056) for 5mins prior to loading and are cultured for 5 hours before stimulation. Cells were cultured using standard conditions for cell culture (5% CO_2_ and 37°C) and maintained using an incubation chamber during imaging. TNF--α (PMC3014_3671982503) was diluted in Fluorobrite DMEM media (A1896701) with 2x glutamax (35050061), pen/strep (15140-122) and FBS (16140071) for stimulation of NIH 3T3 cells. Vials of stimulation media was pressurized at 5psi, kept on ice, and connected to the chip via microbore tubing (PEEK, Vici). The microfluidic device is mounted on the microscope. Knockout p65-/- mouse 3T3 fibroblasts were engineered with p65-DsRed under the native p65 promotor (Tay et al., 2010) and a minimum fluorescence clone was selected to represent endogenous expression of NF-κB and the pathway dynamics. A ubiquitin-promotor driven H2B-GFP cassette provides a nuclear marker for image processing.

We image using Nikon eclipse ti2 microscope to capture both phase and fluorescence images of cultured cells at a 20x magnification. We use a Hamamatsu ORCA-Flash4.0 V3 Digital CMOS Camera (C13440) to capture an image every 5mins for the duration of the experiment. Custom Matlab scripts were used for image processing.

### Mathematical Modeling

See Supplementary Information for an extended discussion of modeling and machine learning methods

## Supplementary Information

### Data processing

Predicting the activation of cells using microscopy images can be formulated as an object classification problem. Data processing details are as follows: we first use min-max normalization to scale the pixel intensity into the range of [0, 1]. Then, we crop each cell as a 64 × 64 image patch centered around the cell nucleus. We split the dataset into 10 folds and set up cross-validation experiment to evaluate our model. In each run, eight folds are used for training, one-fold is used for validation, and the last fold is used for evaluation.

### CNN Model description

We construct a Convolutional Neural Network (CNN) as our predictive model, which follows the similar architecture of LeNet(Lecun *et al.*, 1998). Supplemental Figure 1 shows the model schematic and functionality of each layer. There are two convolutional layers each followed by max-pooling layers. After flattening the output of the second max-pooling layer, two fully connected layers are applied with tanh activation function. Both dense layers are followed by dropout layers with a dropout ratio of 0.5 during training. The last layer is a sigmoid function to calculate the probability of two classes. Testing on a subset of cells that have high model confidence scores within our cross-validation sample results in an overall accuracy of 0.876% +/− 0.036 (Supplementary Figure 1b,c). Accuracy versus prediction score is shown in Supplementary Figure 1d,e.

### CNN Training details

Binary cross-entropy loss is calculated using the final output from the model. Since the label distribution in the dataset is not perfectly balanced, a weighted loss is applied based on the label distribution of the training set. We choose Adam optimizer with a learning rate of 0.001 to optimize the weights from the model. Training is performed in batches of 32 training instances and over 200 epochs. The minimum loss on the validation set determines the optimal time to stop the training process.

### Extraction of Texture Features and SVM Model

Texture feature extraction was done during nuclear segmentation and for each tracked single cell. The number of all extracted parameters is 236 and are shown in **Supplemental Table 1**. Texture extraction was done using several custom Matlab scripts. SVM Model was run using cubic kernel with an overall accuracy of 73%. Fitting was done with 10 fold cross validation with 10% dropout on a dataset with [0.005, 0.05, 0.5, 5 ng/mL] TNF input images (n=3456) as well as a dataset with 0.1ng/mL TNF input images (n=2113) prior to stimulation (Supplemental Figure 4).

### Mathematical modeling of the NF-κB pathway

Using a hybrid stochastic deterministic model of the NF-κB pathway, published previously (Tay *et al.*, 2010; Kellogg and Tay, 2015), we simulated 1000 single cells exposed to 20 TNF concentrations. The hybrid model based on Gillespie algorithm uses verified intrinsic noise in TNF receptor-ligand binding and in transcription of IκBα and A20 which form the main negative-feedback loops leading to oscillations

Simulation was done at 100s time steps with 10hour equilibrium period from initial conditions, 6m of a single TNF pulse, then 2h of evolution. Analysis was done on first peak of resulting cell traces of Nuclear NF-κB. Simulated single cells at t=0 can be found in **Supplementary Table 2** for reference TNF concentrations.

See **Supplementary Table 3** for abbreviations, and **Supplementary Table 4** for parameters. NF-κB and TNFR are distributed in a lognormal distribution with means of 10^5^ and 10^4^ molecules and parameters μ and σ sigma (−1/4, √1/2) and (−1,√2). ODEs for the model are listed below.

### Fast Reactions

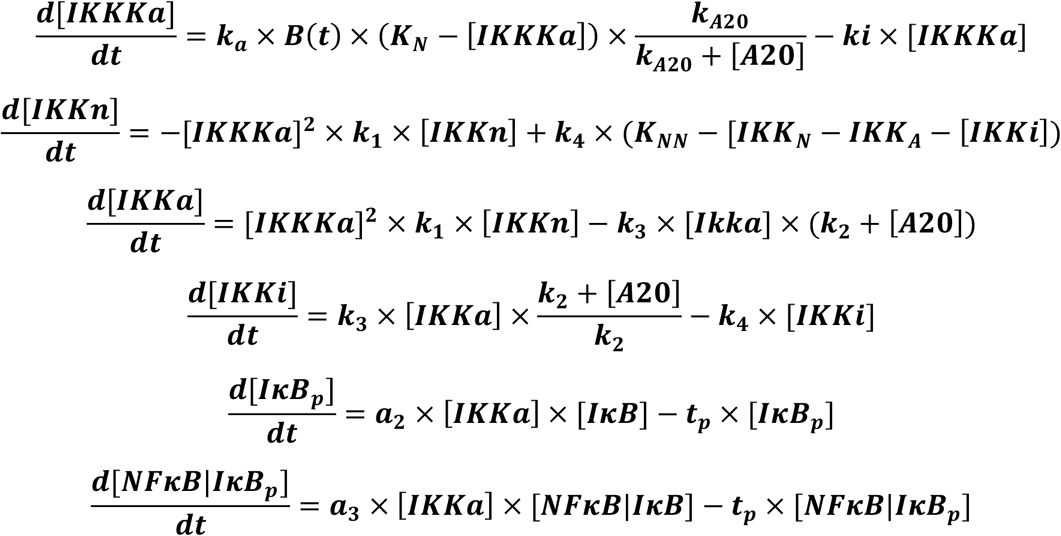

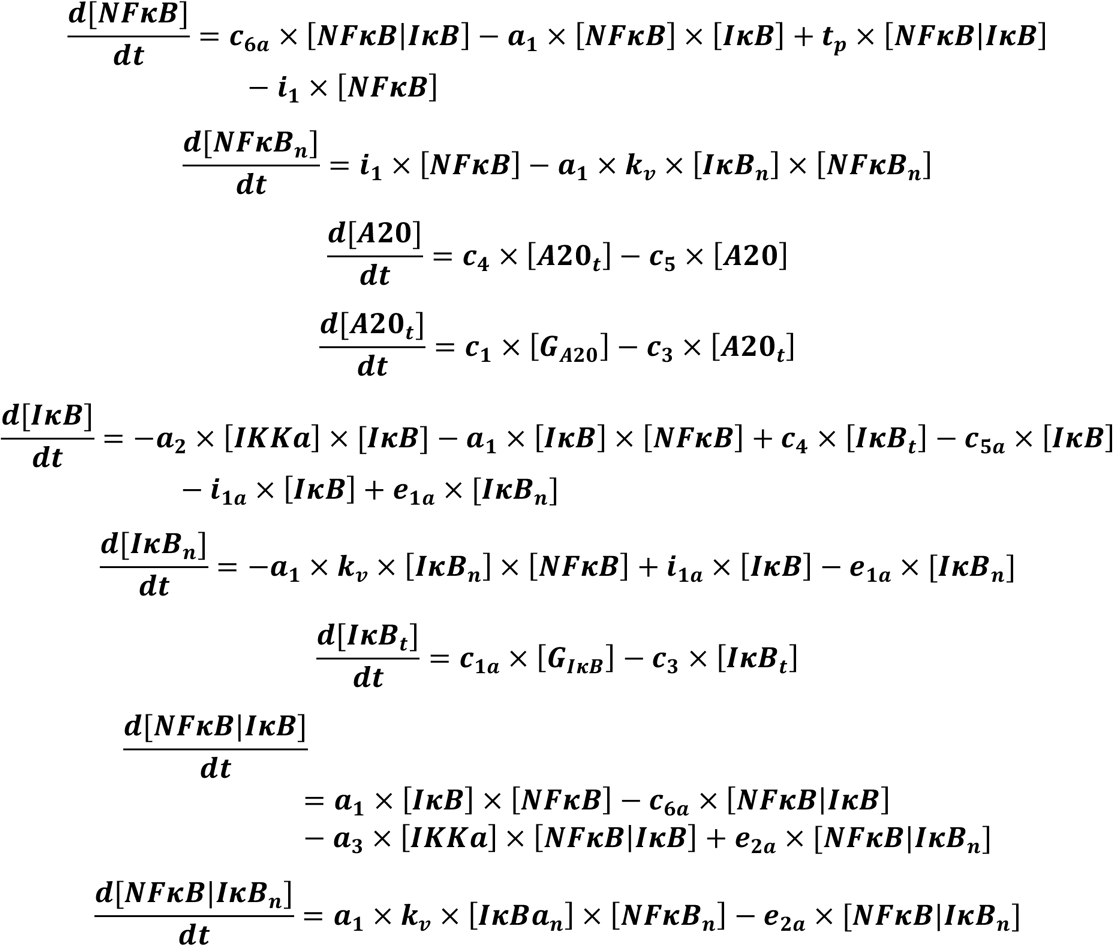

### Reporter is Transcribed Cooperatively

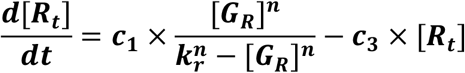

### Slow Reactions

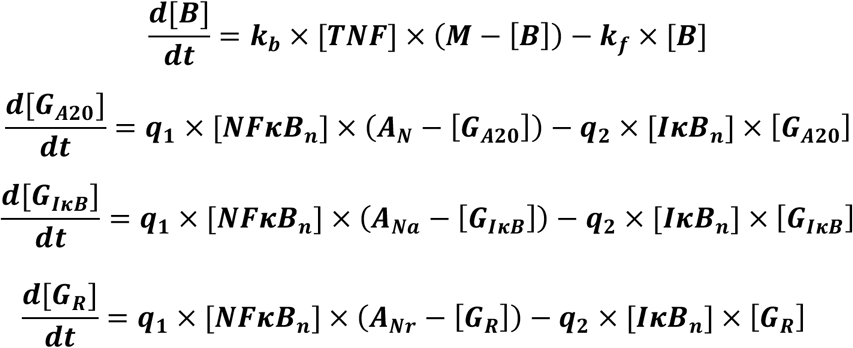

### Stochastic Functions for Receptors, A20, IκB, and reporter genes

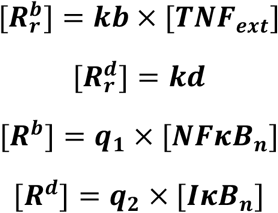

---

**Supplementary Figure 1:**
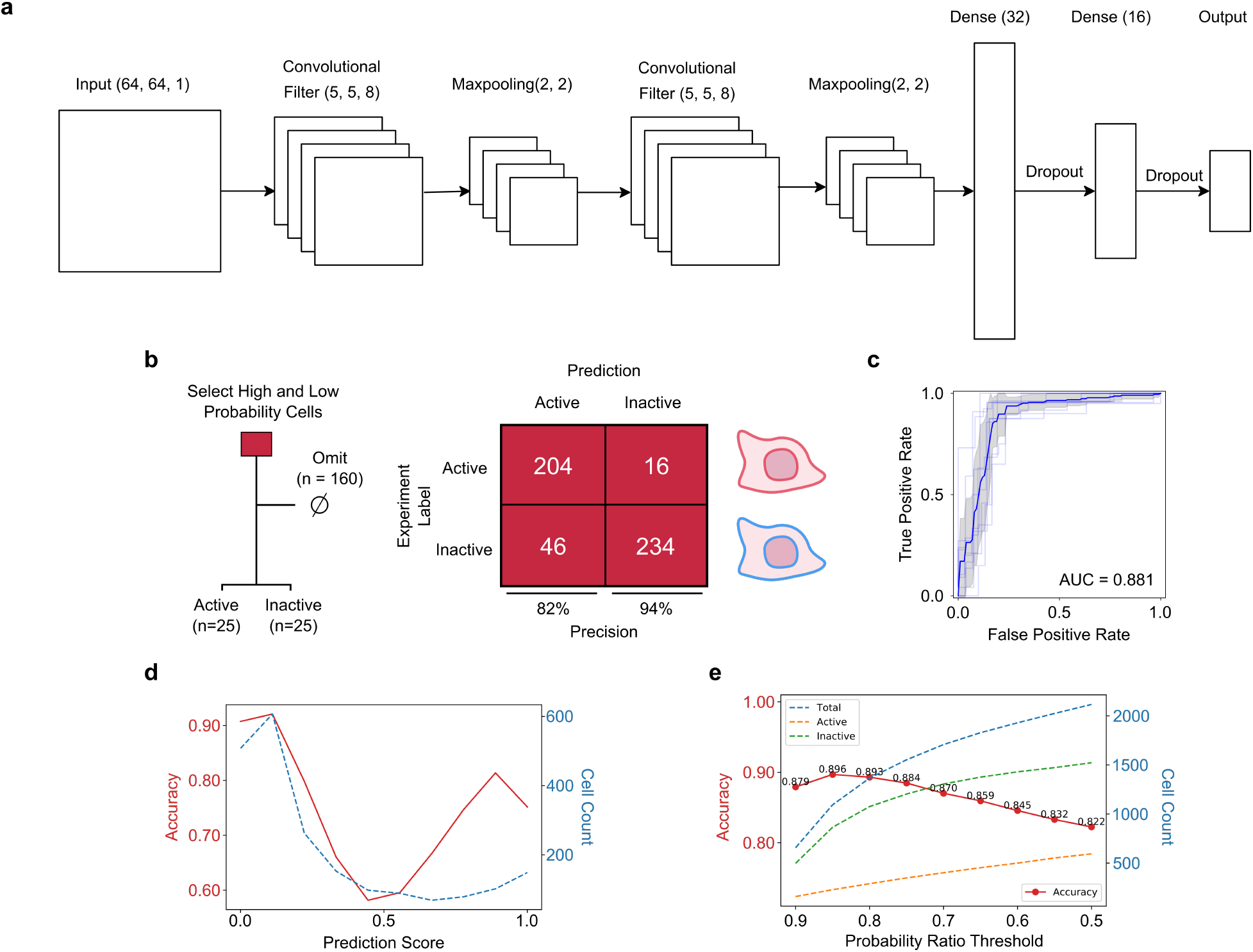
**(a)** Schematic of neural net architecture. Single cell images are passed in as 64×64 pixel images. **(b)** Using a subset of high probability active and inactive cells, we can predict activation and resistance with 82 and 94% precision. **(c)** ROC curve for the highly predictive subset of cells yields an AUC of 0.881. **(d)** There exists a large fraction of cells that will predictably not activate and another fraction that will activate with high fidelity, however, there also exists a subset of cells that remain unpredictable **(e)** Unstimulated fibroblasts have a range of activation probability and on either extreme are highly predictable.

**Supplementary Figure 2:**
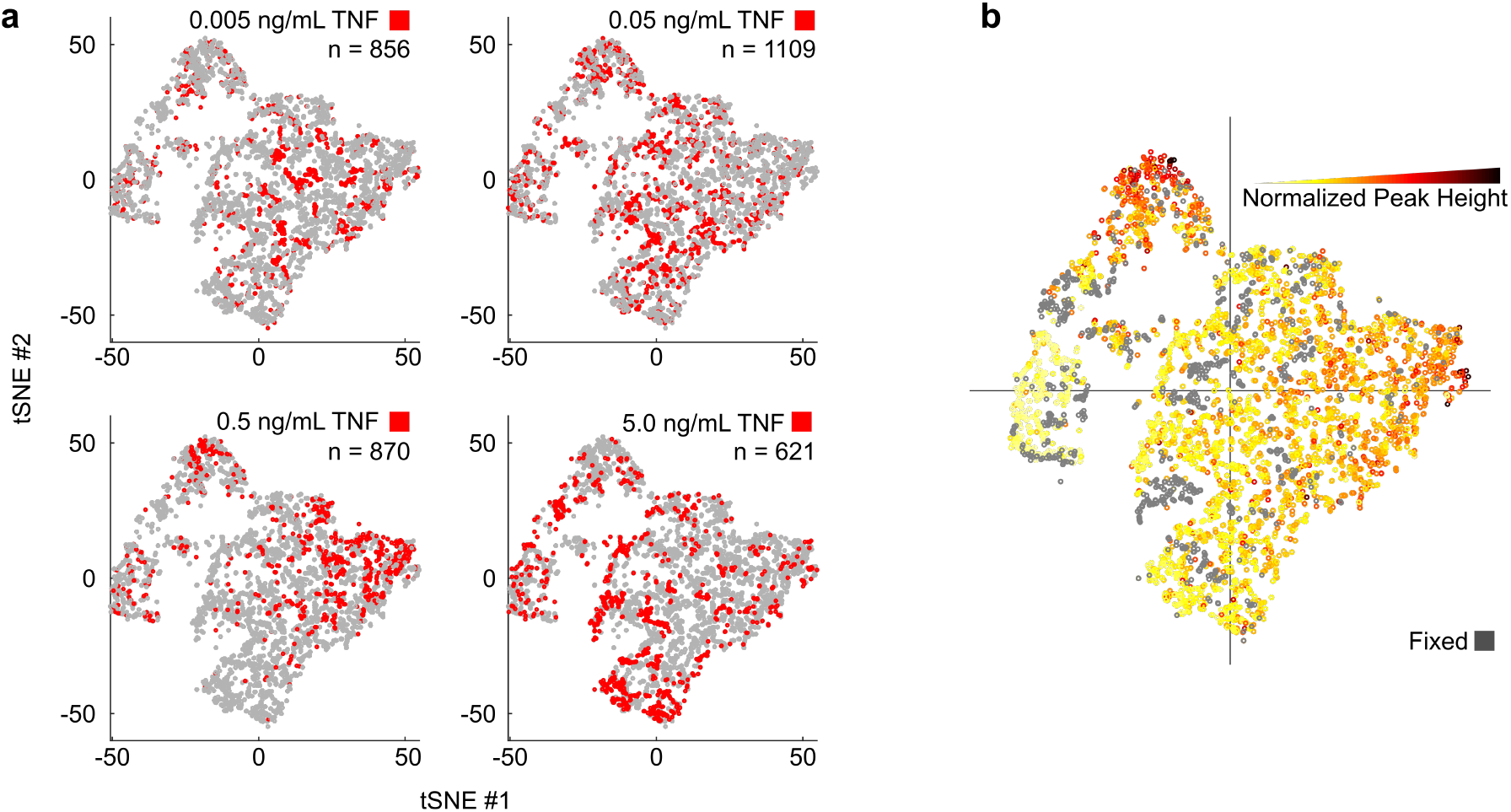
**(a)** tSNE of different TNF dose stimulations (0.005, 0.05, 0.5, 5.0 ng/mL TNF) from t=0 descriptive variables. Cells from a dose are indicated in red. **(b)** tSNE visualization for NF-κB peak heights with overlay of fixed cells.

**Supplementary Figure 3:**
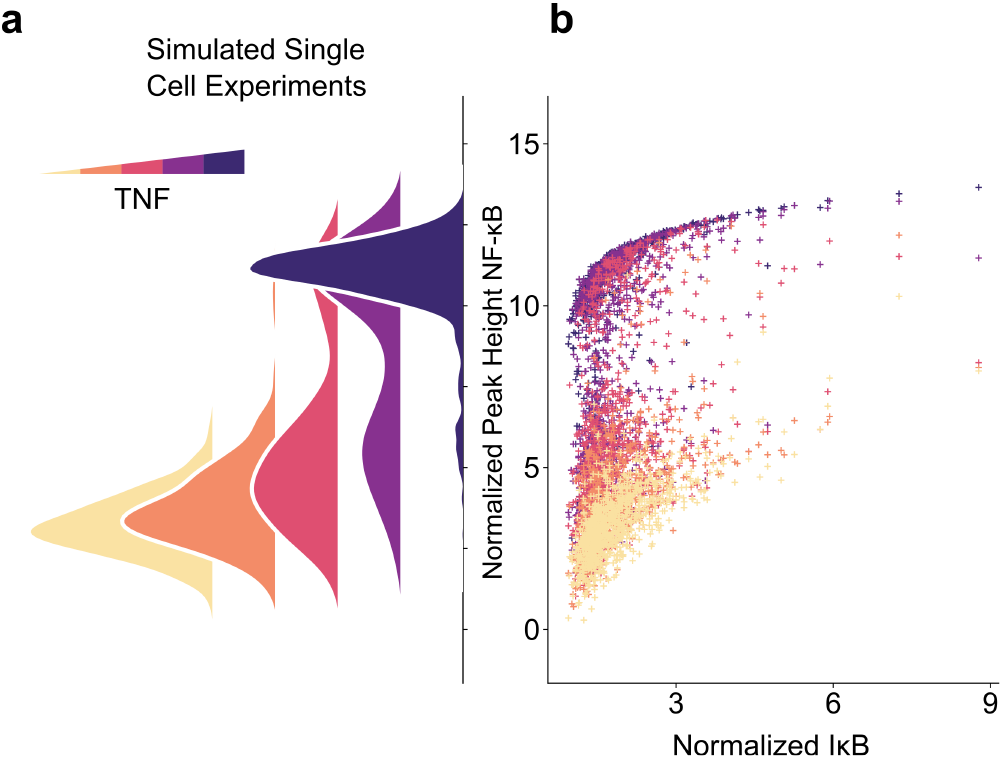
**(a)** Mathematical modeling simulations show single cell nuclear NF-κB peak heights increase with increasing TNF dose. **(b)** High IκB levels in cells require a smaller TNF dose to achieve NF-κB activation.

**Supplementary Figure 4:**
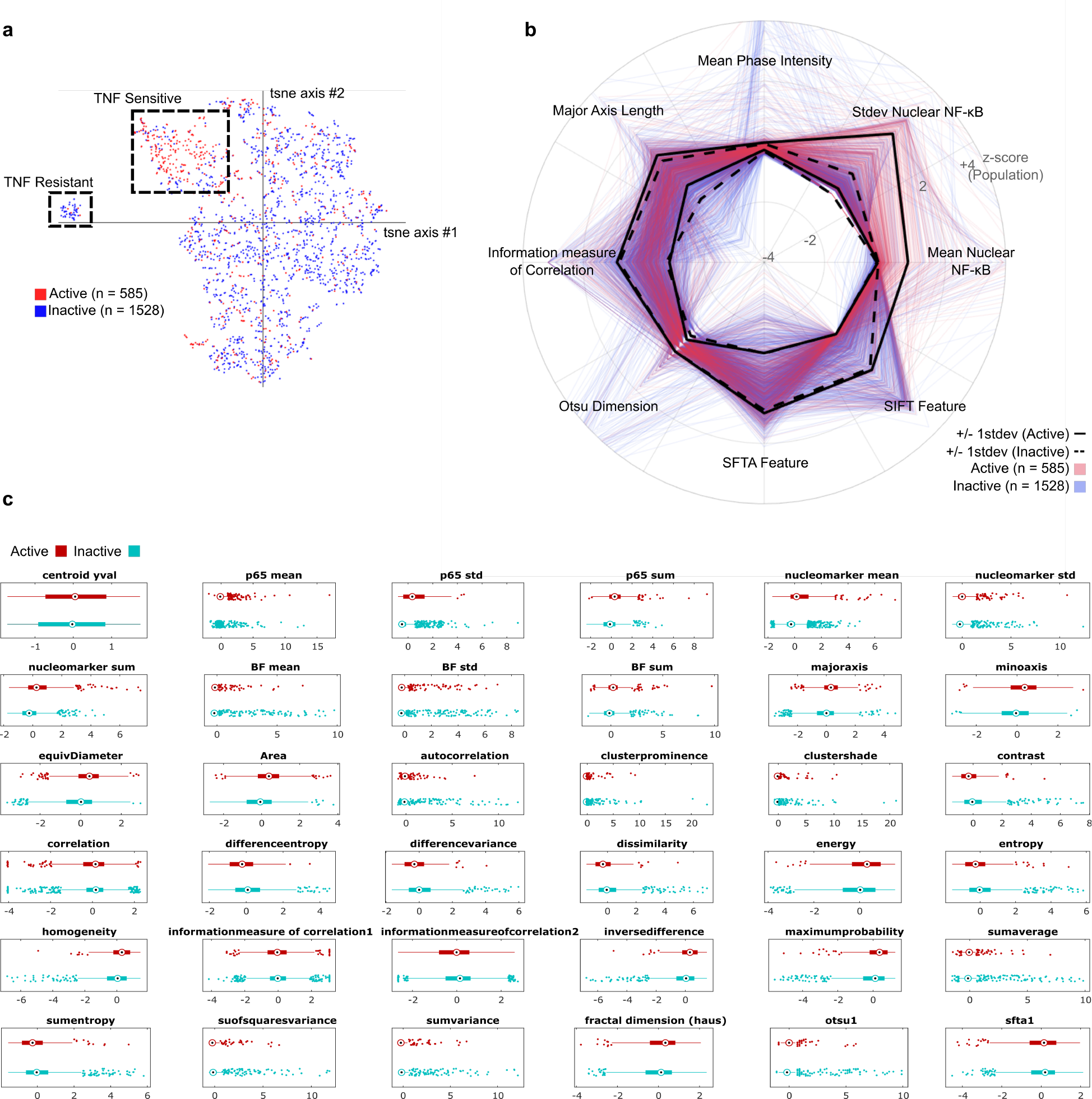
**(a)** tSNE of unstimulated at 0.1ng/mL. **(b)** Highly predictive features for each cell **(c)** Standardized descriptors for unstimulated fibroblasts are shown in boxplots with active cells (red) and inactive cells (blue)

## References

1. Kellogg, R. A., Tian, C., Etzrodt, M. & Tay, S. Cellular Decision Making by Non-Integrative Processing of TLR Inputs. Cell Rep. 19, 125–135 (2017).

2. Ben-David, U. et al. Genetic and transcriptional evolution alters cancer cell line drug response. Nature 560, 325–330 (2018).

3. Chakraborty, A. K. et al. Molecular origin and functional consequences of digital signaling and hysteresis during Ras activation in lymphocytes. Sci. Signal. 2, pt2 (2009).

4. Das, J. et al. Digital signaling and hysteresis characterize ras activation in lymphoid cells. Cell 136, 337–51 (2009).

5. Drayman, N., Patel, P., Vistain, L. & Tay, S. HSV-1 single-cell analysis reveals the activation of anti-viral and developmental programs in distinct sub-populations. Elife 8, (2019).

6. Tay, S. et al. Single-cell NF-kappaB dynamics reveal digital activation and analogue information processing. Nature 466, 267–71 (2010).

7. Avraham, R. et al. Pathogen Cell-to-Cell Variability Drives Heterogeneity in Host Immune Responses. Cell 162, 1309–21 (2015).

8. Lee, M. -C. W. et al. Single-cell analyses of transcriptional heterogeneity during drug tolerance transition in cancer cells by RNA sequencing. Proc. Natl. Acad. Sci. U. S. A. 111, E4726–35 (2014).

9. Otsuki, L. & Brand, A. H. Cell cycle heterogeneity directs the timing of neural stem cell activation from quiescence. Science 360, 99–102 (2018).

10. Sharma, S. V. et al. A Chromatin-Mediated Reversible Drug-Tolerant State in Cancer Cell Subpopulations. Cell 141, 69–80 (2010).

11. Kellogg, R. A., Tian, C., Lipniacki, T., Quake, S. R. & Tay, S. Digital signaling decouples activation probability and population heterogeneity. Elife 4, e08931 (2015).

12. Kellogg, R. A. & Tay, S. Noise Facilitates Transcriptional Control under Dynamic Inputs. Cell 160, 381–392 (2015).

13. Lawrence, T. The nuclear factor NF-kappaB pathway in inflammation. Cold Spring Harb. Perspect. Biol. 1, a001651 (2009).

14. Zhang, Q., Lenardo, M. J. & Baltimore, D. 30 Years of NF-κB: A Blossoming of Relevance to Human Pathobiology. Cell 168, 37–57 (2017).

15. Perkins, N. D. The diverse and complex roles of NF-κB subunits in cancer. Nat. Rev. Cancer 12, 121–132 (2012).

16. Liu, H., Zhang, F., Mishra, S. K., Zhou, S. & Zheng, J. Knowledge-guided fuzzy logic modeling to infer cellular signaling networks from proteomic data. Sci. Rep. 6, 35652 (2016).

17. Cheong, R., Hoffmann, A. & Levchenko, A. Understanding NF-kappaB signaling via mathematical modeling. Mol. Syst. Biol. 4, 192 (2008).

18. Satija, R. & Shalek, A. K. Heterogeneity in immune responses: from populations to single cells. Trends Immunol. 35, 219–229 (2014).

19. Shaffer, S. M. et al. Rare cell variability and drug-induced reprogramming as a mode of cancer drug resistance. Nature 546, 431–435 (2017).

20. Raj, A. & van Oudenaarden, A. Nature, nurture, or chance: stochastic gene expression and its consequences. Cell 135, 216–26 (2008).

21. Kellogg, R. A., Gómez-Sjöberg, R., Leyrat, A. A. & Tay, S. High-throughput microfluidic single-cell analysis pipeline for studies of signaling dynamics. Nat. Protoc. 9, 1713–1726 (2014).

22. Lecun, Y., Bottou, L., Bengio, Y. & Haffner, P. Gradient-based learning applied to document recognition. Proc. IEEE 86, 2278–2324 (1998).

